# Identification of BET Inhibitors (BETi) Against Solitary Fibrous Tumor (SFT) Through High-Throughput Screening (HTS)

**DOI:** 10.1101/2025.03.25.645256

**Authors:** Jose L. Mondaza-Hernandez, David S. Moura, Yi Li, Jesus L. Marti, Paulino Gomez-Puertas, John T. Nguyen, Shuguang Wei, Bruce A. Posner, Clark A. Meyer, Leonidas Bleris, Javier Martin-Broto, Heather N. Hayenga

## Abstract

Cancers, especially fusion oncoprotein (FO)-driven hematological cancers and sarcomas, often develop from a low number of key mutations. Solitary Fibrous Tumor (SFT) is a rare mesenchymal tumor driven by the NAB2-STAT6 oncofusion gene. Currently, the treatment options for SFT remain limited, with anti-angiogenic drugs providing only partial responses and an average survival of two years. To address this challenge, we constructed SFT cell models harboring specific NAB2-STAT6 fusion transcripts using the CRISPR (Clustered Regularly Interspaced Short Palindromic Repeats) technology. High-throughput drug screens demonstrated that the BET inhibitor Mivebresib can differentially reduce proliferation in SFT cell models. Subsequently, BET inhibitors Mivebresib and BMS-986158 efficiently reduced tumor growth in an SFT patient-derived xenograft (PDX) animal model. Furthermore, our data showed that NAB2-STAT6 fusions may lead to higher levels of DNA damage in SFTs. Consequently, combining BET inhibitors with PARP (Poly (ADP-ribose) polymerase) or ATR inhibitors significantly enhanced anti-proliferative effects in SFT cells. Taken together, our study established BET inhibitors Mivebresib and BMS-986158 as promising anti-SFT agents.

**Significance:** New therapies are a clinical need for patients with Solitary Fibrous Tumor. We demonstrated that BET inhibitors are highly active in the preclinical setting for the treatment of this sarcoma entity.

## Introduction

Cancers can result from a relatively low number of mutations developed in key signaling pathways that appear to drive tumorigenesis. Indeed, a study quantifying the tumor mutational burden (TMB) in over 100 tumor types found a median TMB of only 3.6 mutations/Mb^1^. In addition to genomic alterations such as SNPs, insertions, deletions, and copy number alterations, fusion genes also play a critical role in oncogenesis. Tumors with fusion gene events tend to have an even lower overall mutation burden, suggesting that the fusion drives tumorgenicity^2^. A systematic study across several cancer types found that fusions drive the development of 16.5% of cancer cases and are the sole driver in more than 1% of them. Most oncogenic fusion genes are found in hematological cancers, sarcomas, and prostate cancer^3,4^. Elucidating fusion genes in these cancers has led to fusion-specific drugs^5,6^, immunogenic peptides, and immune therapy.

Solitary fibrous tumor (SFT) is a mesenchymal tumor that demonstrates fibroblastic differentiation and may arise anywhere in the body. In 2013, it was discovered that all solitary fibrous tumors have a version of a hallmark intrachromosomal fusion gene between NAB2 and STAT6 on chromosome 12^7,8^. Since then, at least 6 distinct fusion types that account for the observed pathologic variation and tumor aggressiveness have been identified in SFTs, *NAB2_exon6_::STAT6_exon16/17_* and *NAB2_exon4_::STAT6_exon2_* being the most frequent variants^9^. Previous studies suggest that NAB2-STAT6 is the oncogenic driver; otherwise, the TMB in SFT patients has 0 mutations/Mb^7,9,10^. Yet even though this single fusion gene drives tumorigenicity, therapeutic options and, most importantly, targeted studies and clinical trials are lacking. Surgery and/or radiation is the first line of treatment against this tumor; however, for many, this becomes challenging as cancer can travel to inoperable areas or reoccur in locations already irradiated. Anti-angiogenic drugs developed to treat other cancers, including kidney, ovarian, colorectal, lung, and brain, are the best therapeutic options for advanced SFT^11–14^. However, none of the currently available systemic therapies enable complete remission, with the best response being a partial response or stable disease for several months. The average survival rate of patients on the chemotherapies available is 2 years^15^. One of the major bottlenecks in the SFT research field is the lack of *in vitro* and *in vivo* disease models. In this study, we used CRISPR (<u>C</u>lustered <u>R</u>egularly Interspaced <u>S</u>hort <u>P</u>alindromic <u>R</u>epeats) based genome editing to engineer SFT cell models with specific NAB2-STAT6 gene fusions^5,6^, which were subsequently applied to a high-throughput screening (HTS) platform. We identified compounds that selectively disrupt the oncogenic NAB2-STAT6 fusion-driven signaling, which could later be used as systemic therapeutic agents for SFT. Finally, the anti-tumor efficacies of candidate compounds were validated using an SFT patient-derived xenograft (PDX) mouse model.

## Results

### Primary High-Throughput Screening (HTS) using the SFT NS-poly cells

Using the CRISPR/spCas9 system, we first generated an engineered SFT cell line for the NAB2_exon6_-STAT6_exon17_ fusion type in HCT116 cells (NS-poly). Compared to prior SFT cell models^10^, NS-poly cells preserve all original *NAB2-STAT6* gene fusion information, including endogenous NAB2 promoters, 5’-UTRs (5’-untranslated region) of NAB2, and 3’-UTRs of STAT6. Next, we performed a primary HTS assay using the NS-poly cells against the FDA-approved and experimental drug collection (∼2,600 compounds)^16–23^. Briefly, NS-poly cells were seeded into 384-well assay plates and treated with the annotated chemical library at a final concentration of 5 µM. The assay plates were then processed as described in Materials and methods/ High-Throughput Screening (HTS). For quantitative analysis, the fractions of dead/dying cells (DRAQ7 cell count/CellTracker Deep Red cell count) were calculated for each well at all time points, which were used to plot the dose-response curves (e.g., Supplementary Figure S1a for Paclitaxel in NS-poly cells). Subsequently, we computed an area-under-the-curve (AUC) result for the treatment time course (e.g., Supplementary Figure S1b for Paclitaxel in NS-poly cells). We used two normalization methods to assess the activity of library compounds: a test population-based method and a controls-based approach. For the former, we calculated a robust mean and standard deviation for the test population (library compound containing wells) and scaled compound activities to DMSO (arbitrarily set to 0% effect). For the latter normalization, we calculated the robust mean and standard deviation for the controls (vehicle control: DMSO alone and positive control: paclitaxel) and scaled compound activity (observed – DMSO mean) to the difference between the DMSO and paclitaxel controls. The activities observed for DMSO were set to 0% effect, and the activities of library compounds were scaled accordingly. Finally, to identify the hit compounds that can efficiently induce cell death in NS-poly cells, the robust Z-scores for both (a) normalized activities at the final timepoint (72 hours, named as Final Timepoint effects) and (b) normalized AUCs (named as AUC effects) were calculated^24^. Only compounds with robust Z-scores less than -3 were selected for the following secondary assays and counter screens. 247 compounds were identified using the Final Timepoint effects (Supplementary Table S1), and 232 were determined using the AUC effects (Supplementary Table S2).

### Secondary High-Throughput Screening (HTS) using patient-derived Moffitt-ns and immortalized lung fibroblast cells

In our selection process for primary hits for secondary HTS assays, we first identified compounds that passed the primary screening criteria (robust Z-score < -3) and exerted suppression effects of >70% for both the 72 hours endpoint (final timepoint effects) and the AUC measurements (AUC effects). In addition to the hits identified by the primary HTS, we included systemic agents currently used in or related to treating SFT patients (e.g., receptor tyrosine kinases inhibitor Sunitinib). In total, 104 compounds (Supplementary Table S3, 93 from the primary screen and 11 from the clinical applications) were procured from either Selleck Chemicals (60) or NIH (44).

Given the colorectal cancer background, our HCT116-based NS-poly cell model may harbor additional oncogenic drivers other than the NAB2-STAT6 fusion, which could muddle the interpretation of HTS results. Therefore, in parallel, we established a primary cell line from an SFT patient at Moffitt Cancer Center (named Moffitt-ns, fusion type: NAB2_exon5_-STAT6_exon16_)^6^. Finally, to match the fibrous background of Moffitt-ns, an hTERT-immortalized human lung fibroblast cell line (Lf) was used as the negative control. The same screening platform (dual dyes: CellTracker Deep Red and DRAQ7) from the primary screen was adopted for the secondary screen, except that: (a) Four doses (50 nM, 200 nM, 650 nM, and 2 µM for compounds from Selleck Chemicals; 125 nM, 500 nM, 1.6 µM, and 5 µM for compounds from the NIH Clinical Collection) were tested; and (b) an additional cell viability assay (Promega, CellTiter-Glo Luminescent Cell Viability assay) was performed at the final timepoint (CTG effects).

To identify candidate compounds that can selectively, efficiently, and safely suppress the growth of Moffitt-ns cells, we applied three filtering conditions: 1) Selectivity: The differences in cell suppression between Moffitt-ns and control Lf are > 40% for at least two doses; 2) Efficacy: Cell suppression at the highest dose should be > 50% in Moffitt-ns cells; and, 3) Off-target toxicity: Cell suppression at the highest dose should be < 50% in the control Lf cells. As shown in Figure 1a and Supplementary Tables S4 and S5, using these filtering conditions, we identified 3 candidates (Mivebresib, Zinc Pyrithione, and TAK-901) for the CTG effects, and 11 candidates (Mivebresib, Zinc Pyrithione, Staurosporine, Digoxin, Lanatoside C, Proscillaridin A, Disulfiram, Doxorubicin hydrochloride, Colchicine, Vinorelbine, and SW197775) for the final timepoint effects (Supplementary Tables S6 and S7). Similarly, for the AUC effects (Supplementary Tables S8 and S9), 8 hits were identified (Zinc Pyrithione, Staurosporine, Digoxin, Lanatoside C, Proscillaridin A, Doxorubicin hydrochloride, Colchicine, and SW197775).

**Figure 1.**
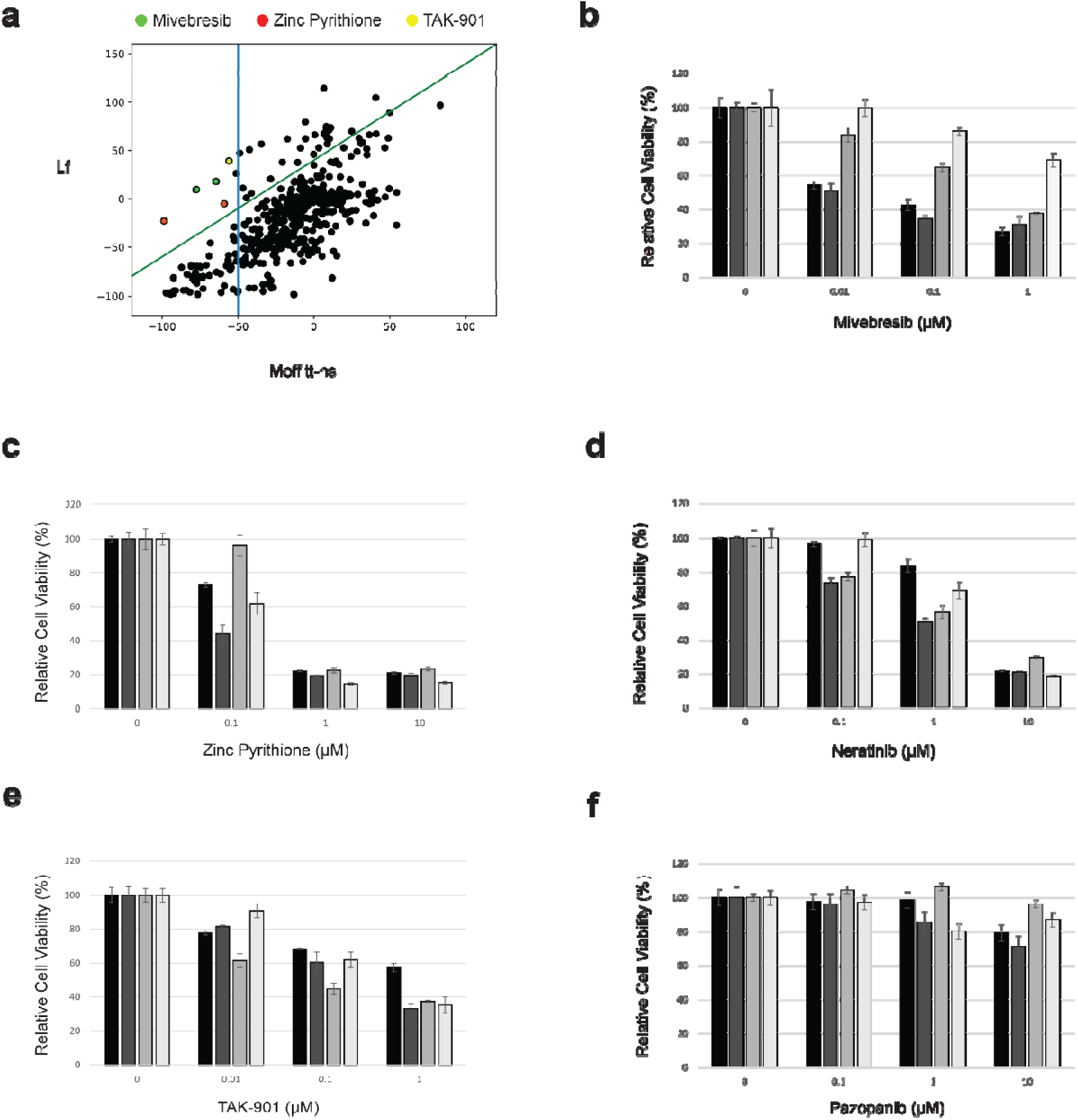
Identification of Miverbresib and Zinc Pyrithione through High-Throughput Screenings (HTS) against Solitary Fibrous Tumor (SFT). (**a**) Secondary high-throughput screening (HTS) was performed using the Moffitt-ns and immortalized lung fibroblast cells. The CTG effects of both cell lines under all four doses were plotted. The green line indicated the differences in suppression effects between Moffitt-ns and Lf > 40%. The blue line indicated that the suppression effects at the highest dose should be > 50% in Moffitt-ns cells. 3 compounds were shown (Mivebresib: green, Zinc Pyrithione: red, and TAK-901: yellow). (**b-f**) Confirmatory MTS assays were performed in two primary SFT cells (INT-SFT and IEC139) and two control LMS cells (SKUT-1 and CP0024). Only Mivebresib showed efficient and selective cell proliferation-suppressing effects in SFT cells. (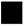): INT-SFT; (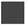): IEC139; (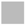): SKUT-1; (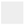): CP0024.

### *In vitro* efficacy testing of Mivebresib using additional primary SFT cells

To determine candidate compounds for further efficacy studies, we focused on compounds that suppress the growth of SFT cancer cells. Thus, more weight was given to the CTG and final timepoint effects, and subsequently, two hits (Mivebresib, Zinc Pyrithione) were identified as present in both analyses (Supplementary Tables S4-S7). It is worth noting that Zinc Pyrithione was also identified in the AUC effects analysis (Supplementary Table S9). For controls, we included three compounds from our secondary screening that did not pass the filtering conditions (Neratinib, TAK-901, and Pazopanib).

Our primary and secondary HTS assays were performed using specific NAB2-STAT6 fusion types (NAB2_exon6_-STAT6_exon17_ for the primary screen and NAB2_exon5_-STAT6_exon16_ for the secondary screen). To further test the robustness of our candidate compounds, we performed *in vitro* confirmatory efficacy testing (MTS Cell Viability assay) in two additional SFT patient-derived primary cell lines (INT-SFT with a fusion type: NAB2_exon6_-NAB2_intron6_-STAT6_exon16_; and IEC139 with a fusion type: NAB2_exon6_-STAT6_exon16_) consisting of different NAB2-STAT6 fusion types^25^. As shown in Figure 1b, Mivebresib efficiently suppressed the cell growth in both SFT cell lines (IC_50_: 8.94 nM for INT-SFT, and 7.71 nM for IEC139), while much less so in two Leiomyosarcoma (LMS) cell lines (IC_50_: 346.90 nM for SKUT-1, and 4.17 µM for CP0024). In comparison, other tested compounds either showed poor differential effects between SFT and LMS cell lines (Figures 1c, 1d, and 1e for Zinc Pyrithione, Neratinib, and TAK-901, respectively) or no clinically meaningful effects in all cell lines (Figure 1f for Pazopanib). As an example (Table 1), the IC_50_ values for Neratinib were 3.61 µM, 0.88 µM, 1.59 µM, and 1.34 µM for INT-SFT, IEC139, SKUT-1, and CP0024 cells, respectively. These results indicated that Mivebresib can efficiently and selectively suppress SFT cells *in vitro*.

**Table 1.**
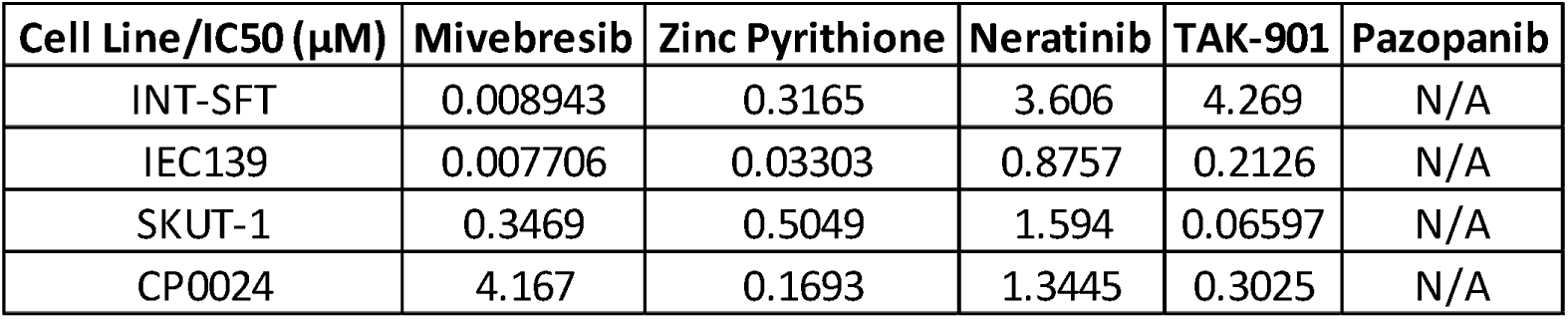
IC50 values for candidate compounds in SFT and LMS cell models. N/A: not applicable.

| **Cell Line/IC50 (μM)** | **Mivebresib** | **Zinc Pyrithione** | **Neratinib** | **TAK-901** | **Pazopanib** |
| --- | --- | --- | --- | --- | --- |
| INT-SFT | 0.008943 | 0.3165 | 3.606 | 4.269 | N/A |
| IEC139 | 0.007706 | 0.03303 | 0.8757 | 0.2126 | N/A |
| SKUT-1 | 0.3469 | 0.5049 | 1.594 | 0.06597 | N/A |
| CP0024 | 4.167 | 0.1693 | 1.3445 | 0.3025 | N/A |

### *In vitro* efficacy testing of additional BETis

Although Mivebresib, a pan BETi, showed promising efficacy and specificity in SFT cells (Figure 1), it is worth noting that in previous phase I clinical trials, some side effects, including thrombocytopenia and anemia, were observed^26,27^. Therefore, we next evaluated six additional BETis (Supplementary Table S10, BMS-986158^28^, Pelabresib^29^, ABBV-744^30^, PLX51107^31^, GSK778^32^, and GSK046^32^) with different targeting selectivity for their *in vitro* efficacy and specificity against SFTs. As shown in Figures 2a and 2b, BD1- and BD2-selective BETis (GSK778 for BD1, ABBV-7444 and GSK046 for BD2), as well as pan-BETis Pelabresib and PLX51107, did not significantly decrease cell viability in both INT-SFT and IEC139 cells. In contrast, pan-BETi BMS-986158 (50 nM, 72 hours treatment) potently induced cell apoptosis and necrosis (45.9% for INT-SFT, 34.3% for IEC139) using Annexin V/PI staining-based flow cytometry assay. Similarly, the MTS cell viability assay showed that while BMS-986158 effectively suppressed the cell proliferation in INT-SFT and IEC139 cells (IC_50_ values: 6.23 nM for INT-SFT and 28.8 nM for IEC139), it showed no anti-proliferative effects in LMS, CP0024, cells (Figure 2c). Interestingly, unlike Mivebresib, BMS-986158 also potently suppressed the growth of another LMS cell line SKUT-1 (IC_50_: 3.38 nM). Taken together, these results showed that BMS-986158 demonstrated high efficiency in suppressing the growth of SFT cells *in vitro* but lower SFT specificity compared to Mivebresib.

**Figure 2.**
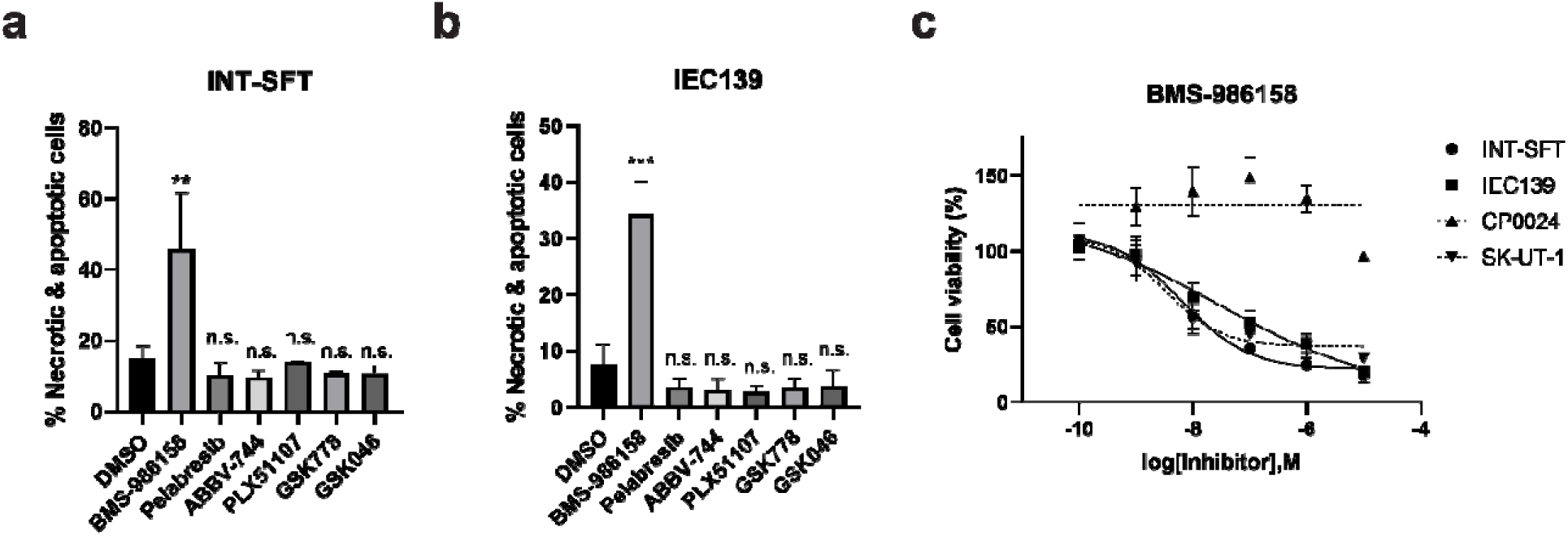
*In vitro* evaluation of activities and specificities of additional BET inhibitors in SFT cells. **a)** *In vitro* evaluation of activities of additional BET inhibitors (BMS-986158, Pelabresib, ABBV-744, PLX51107, GSK778, and GSK046) in INT-SFT cells using Flow cytometry-based apoptosis analysis. **b)** *In vitro* evaluation of activities of additional BET inhibitors (BMS-986158, Pelabresib, ABBV-744, PLX51107, GSK778, and GSK046) in IEC139 cells using Flow cytometry-based apoptosis analysis. For both **a)** and **b)**, cells were treated with candidate chemicals (50 nM) for 72 hours, and the drug activities were expressed as the percentages of non-viable cell subpopulation to the total cell population. For statistical analysis, two-tailed t-tests were conducted. ** denotes p<0.01, *** denotes p<0.001, n.s. denotes no significant difference. **c)** Dose-response curves for BMS-986158 at 72 hours in primary SFT cells (INT-SFT and IEC139) and control leiomyosarcoma (CP0024 and SK-UT-1) cells using MTS cell viability assays. Cell viability was normalized to untreated conditions (n=4).

### BETis Mivebresib and BMS-986158 induced DNA breaks and G1 cell cycle arrest in SFT cells

To uncover the molecular mechanisms of the anti-proliferation effects of Mivebresib and BMS-986158 in SFT cells, we noted that the treatment of these BETis (50 nM, 72 hours) induced the cleavage of PARP-1 (poly(ADP-ribose) polymerase 1, Figure 3a), indicating the activation of apoptosis signaling pathways^33,34^. In addition, the phosphorylation of H2AX at serine 139 (γ-H2AX), a recognized marker of double-strand DNA breaks (DSBs) and replication stress^35,36^ was observed (Figure 3a). More specifically, the treatment of Mivebresib led to a 19.6- and 22.5-fold increase in γ-H2AX levels in INT-SFT and IEC139 cells, respectively, while the treatment of BMS-986158 resulted in a 49.1- and 31.6-fold increase in the same cell lines.

**Figure 3.**
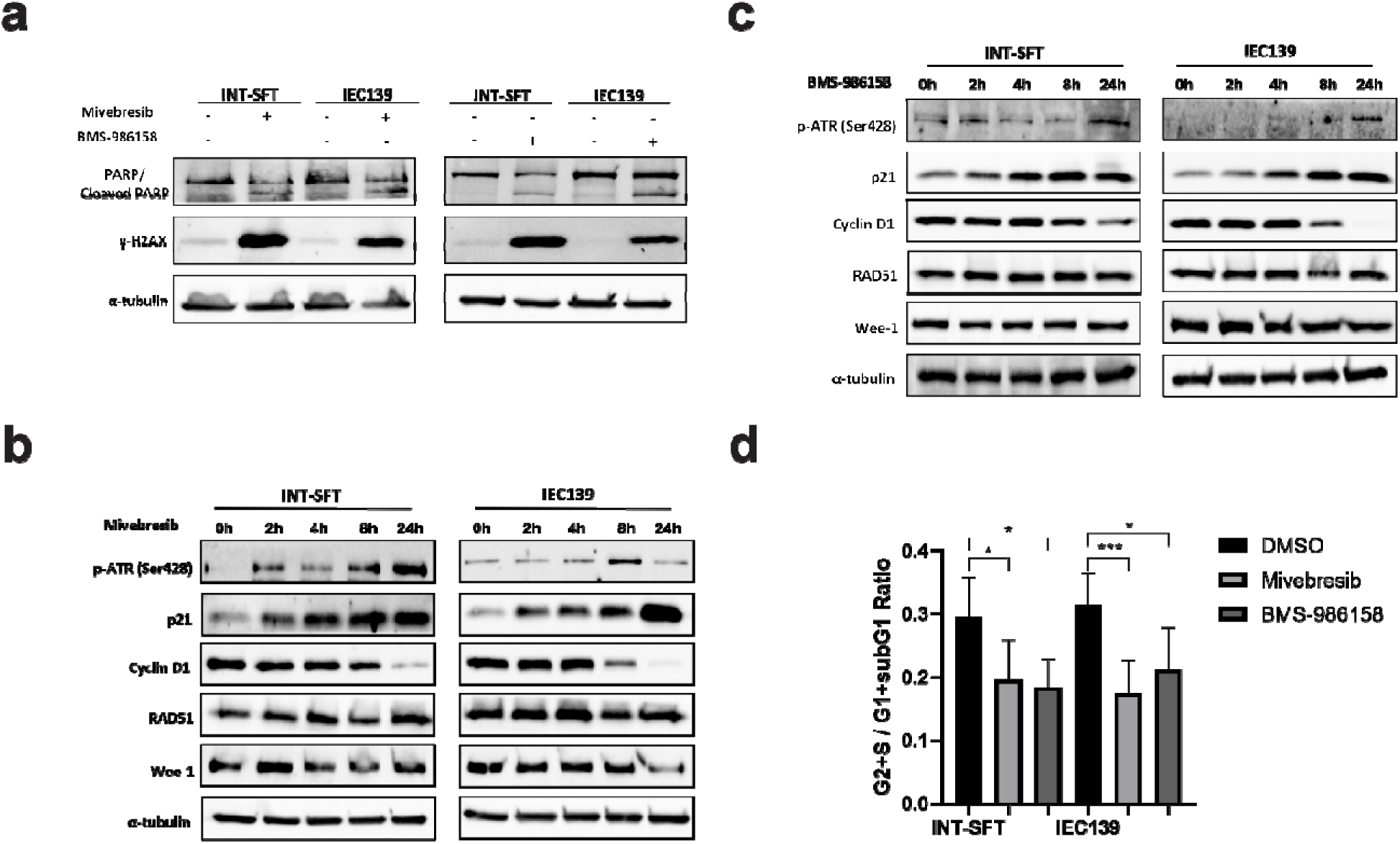
BET inhibitors (BETi) Mivebresib and BMS-986158 exerted anti-proliferative effects in SFTs via cell cycle and DNA damage response pathways. **a)** Protein levels of apoptosis marker (cleaved PARP-1) and DSBs (double strand-breaks) marker (γ-H2AX) increased in SFT cells after 72-hour treatment with 50 nM of Mivebresib or BMS-986158. **b)** Protein levels of phosphorylated ATR at serine 428 (p-ATR, Ser428), p21, Cyclin D1, RAD51, and Wee-1 in SFT cells (INT-SFT and IEC139) during a 24-hour treatment with 50 nM Mivebresib. The α-tubulin was used as a loading control. **c)** Protein levels of phosphorylated ATR at serine 428 (p-ATR, Ser428), p21, Cyclin D1, RAD51, and Wee-1 in SFT cells (INT-SFT and IEC139) during a 24-hour treatment with 50 nM BMS-986158. The α-tubulin was used as a loading control. d) Ratios representing proliferating (G2 and S) vs non-proliferating (sub-G1 and G1) cells upon Mivebresib or BMS-986158 treatment. Bar plots represent mean values with standard deviations. For statistical analysis, two-tailed t-tests were conducted. * denotes p < 0.05; *** denotes p < 0.001.

We further analyzed additional protein markers associated with DNA damage response (DDR) pathways or cell cycle regulation (0 to 24 hours post-BETi treatment). Interestingly, upregulation of ATR phosphorylation at serine 428 was observed for both BETis in SFT cell lines (Figure 3b for Mivebresib, Figure 3c for BMS-986158), suggesting that single-strand DNA (SSD) break repair mechanisms were also activated in response to BETi treatments^37^. Specifically, for Mivebresib, ATR phosphorylation peaked at 24 hours in INT-SFT cells (a 5.5-fold increase compared to baseline) and at 8 hours in IEC139 cells (a 2.9- fold increase compared to baseline). Similarly, for BMS-986158, both cell lines exhibited peak ATR phosphorylation at 24 hours (a 2.1-fold increase for INT-SFT and a 3.6-fold increase for IEC139 compared to baseline).

Additionally, elevated levels of p21^38^ and reduced levels of Cyclin D1^39,40^ were observed upon the drug treatments, both of which implied the activation of the DDR pathway and potential G1 cell cycle arrest. Specifically, 24 hours after the treatment of Mivebresib, the expression levels of p21 increased by 8.0- and 6.1-fold in INT-SFT and IEC139 cells, respectively. Similarly, for BMS-986158, the expression levels of p21 levels increased by 2.8-fold in INT-SFT cells and 29.2-fold in IEC139 cells. Next, the expression levels of cyclin D1 decreased by 72.4% and 98.6% after 24 hours of Mivebresib treatment and by 68.4% and 90.1% after BMS-986158 treatment in INT-SFT and IEC139 cells, respectively (Figure 3b and Figure 3c). Reassuringly, these results were consistent with our propidium iodide-based flow cytometry assays. Briefly, 24 hours after BETis treatments, the ratio of the proliferating and dividing cells (defined as in S and G2 phases) to non-proliferating cells (defined as in G1 phase and sub-G1 phase for INT-SFT cells, see Materials and methods/Apoptosis and Cell Cycle Analysis by Flow Cytometry) was calculated. As shown in Figure 3d, in both SFT cell lines, treatments of Mivebresib and BMS-986158 significantly decreased such ratios, indicating an accumulation of cells in the G1 phase. As an example, in INT-SFT cells, the ratio for cells under the control treatment (0.29) was significantly higher than those under the treatment of either Mivebresib (0.19, p = 0.047) or BMS-986158 (0.18, p = 0.029). Similarly, in IEC139 cells, the DMSO control group demonstrated a higher ratio (0.31) compared to Mivebresib (0.17, p < 0.001) and BMS-986158 (0.21, p = 0.026)-treated groups.

Lastly, we noted that no significant changes were observed in the expression levels of other DNA repair-related proteins, such as RAD51 and Wee1, suggesting that homologous repair mechanisms (RAD51 as the marker^41,42^) were not activated, and checkpoints beyond the G1 phase of the cell cycle (Wee1 as the marker^43^) remained unaffected (Figure 3b and Figure 3c). Taken together, these findings showed that BETis Mivebresib and BMS-986158 may suppress the proliferation of SFT cells via the induction of DNA damage and G1 cell cycle arrest.

### Combinatorial effects between BETis and PARP/ATR inhibitors in SFT cells

Our data showed that BETis Mivebresib and BMS-986158 induced both double-strand DNA break repair (induction of the cleavage of PARP-1) and single-strand DNA break repair (upregulation of ATR phosphorylation) mechanisms in SFT cells. Next, to determine the combinatorial effects between BETis (Mivebresib: 50 nM, BMS-986158: 50 nM) and DNA damage response pathway inhibitors (DDRi): PARP inhibitor Rucaparib^44^: 10 μM, and ATR inhibitor Berzoertib^45^: 1 μM) in SFTs, two assays were utilized: a flow cytometry-based apoptosis assay and Western blot for DDR-related protein markers.

As shown in Figure 4a, as compared to Mivebresib alone, the combination of Mivebresib and Rucaparib induced significantly more apoptosis in both INT-SFT and IEC139 cells (1.4- and 1.5-fold increases for INT-SFT and IEC139, respectively). These results were consistent with the finding that the combinatorial drug treatment induced higher expression levels of both cleaved PARP-1 and γ-H2AX compared to Mivebresib alone (Figure 4b, 1.4-fold increase for cleaved PARP-1 in INT-SFT, 2.5-fold increase for γ-H2AX in INT-SFT, 1.6-fold increase for cleaved PARP-1 in IEC139, and 2.3-fold increase for γ-H2AX in IEC139). Similarly, the combination of BMS-986158 and Rucaparib also induced significantly more apoptosis in both INT-SFT and IEC139 cells (Figure 4c, 1.5- and 1.5-fold increases for INT-SFT and IEC139, respectively), as well as higher expression levels of both cleaved PARP-1 and γ-H2AX (Figure 4d, 1.6-fold increase for cleaved PARP-1 in INT-SFT, 1.3-fold increase for γ-H2AX in INT-SFT, 2.1-fold increase for cleaved PARP-1 in IEC139, and 2.9-fold increase for γ-H2AX in IEC139).

**Figure 4.**
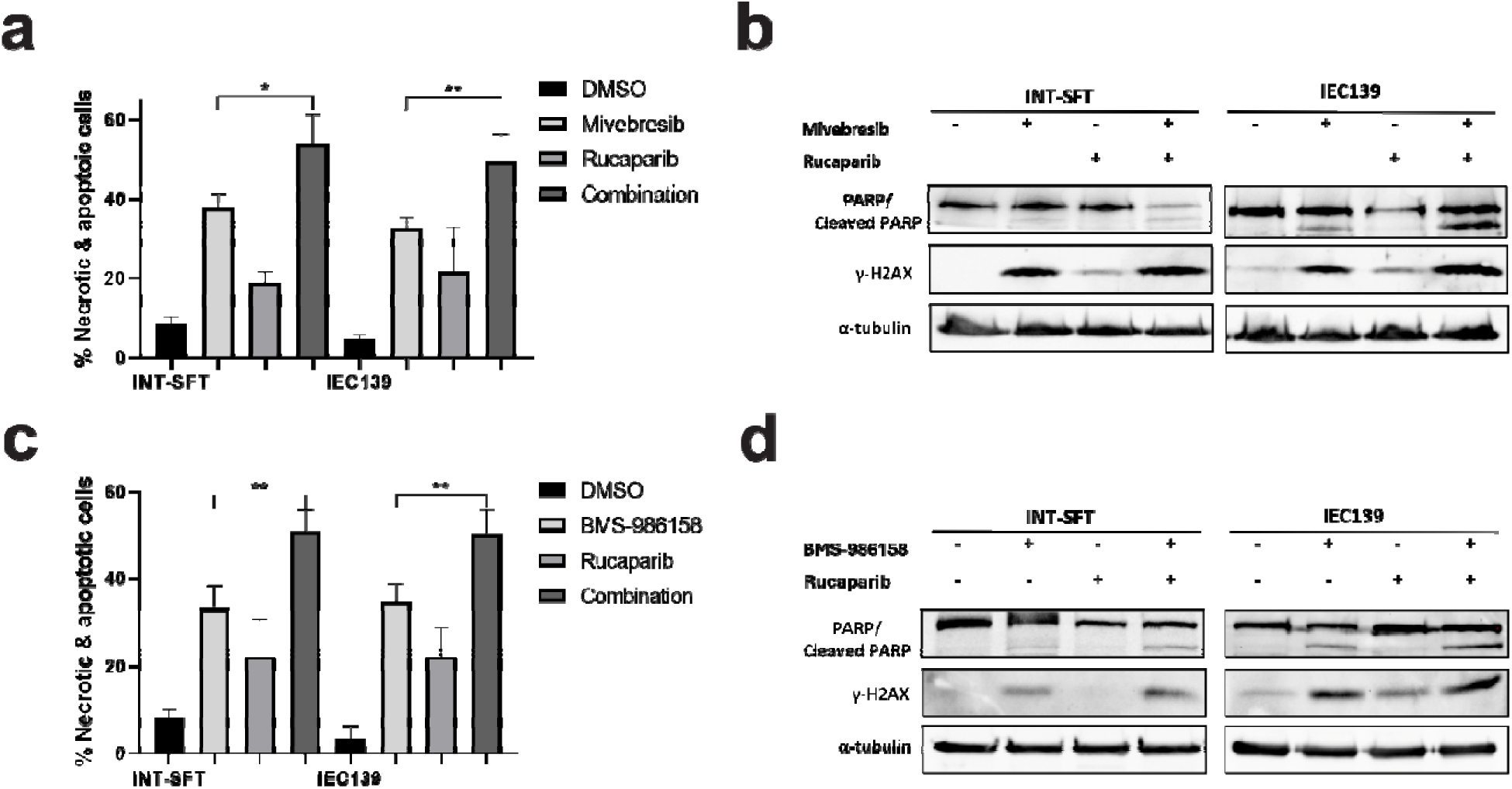
Combinatorial effects between BET inhibitors (Mivebresib and BMS-986158) and PARPi Rucaparib. **a)** Flow cytometry-based apoptosis assays showed that combining Mivebresib and Rucaparib increased apoptotic and necrotic cell populations in INT-SFT and IEC139 cells (n=3). **b)** Western blot assays showed that combining Mivebresib and Rucaparib increased cleaved PARP-1 and γ-H2AX protein levels in INT-SFT and IEC139 cells after 72-hour treatment (n=3). The α-tubulin was used as a loading control. **c)** Flow-cytometry-based apoptosis assays showed that combining BMS-986158 and Rucaparib increased apoptotic and necrotic cell populations in INT-SFT and IEC139 cells (n=3). **d)** Western blot assays showed that combining BMS-986158 and Rucaparib increased cleaved PARP-1 and γ-H2AX protein levels in INT-SFT and IEC139 cells after 72-hour treatment (n=3). The α-tubulin was used as a loading control. For statistical analysis, two-tailed t-tests were conducted. * denotes p < 0.05; ** denotes p < 0.01.

Similar results were observed when combining BETis with ATR inhibitor Berzoertib. As shown in Figure 5a, the combination of Mivebresib and Berzoertib increased cell apoptosis by 1.4- and 1.6-fold compared to Mivebresib alone for INT-SFT and IEC139 cells, respectively (p-values = 0.003 and 0.002, respectively). In addition, the combinatorial drug treatment induced higher expression levels of both cleaved PARP-1 and γ-H2AX compared to Mivebresib alone (Figure 5b, 3.0-fold increase for cleaved PARP-1 in INT-SFT, 1.6-fold increase for γ-H2AX in INT-SFT, 2.1-fold increase for cleaved PARP-1 in IEC139, and 2.0-fold increase for γ-H2AX in IEC139). Similarly, the combination of BMS-986158 and Berzoertib induced more apoptosis in both INT-SFT and IEC139 cells (Figure 5c, 1.5- and 1.7-fold increases for INT-SFT and IEC139, respectively), as well as higher expression levels of both cleaved PARP-1 and γ-H2AX (Figure 5d, 1.6-fold increase for cleaved PARP-1 in INT-SFT, 1.6-fold increase for γ-H2AX in INT-SFT, 1.3-fold increase for cleaved PARP-1 in IEC139, and 2.3-fold increase for γ-H2AX in IEC139). Thus, our results showed that the combinational treatment between our candidate BETis and select PARP/ATR inhibitors exerted higher anti-proliferative effects in SFT cells.

**Figure 5.**
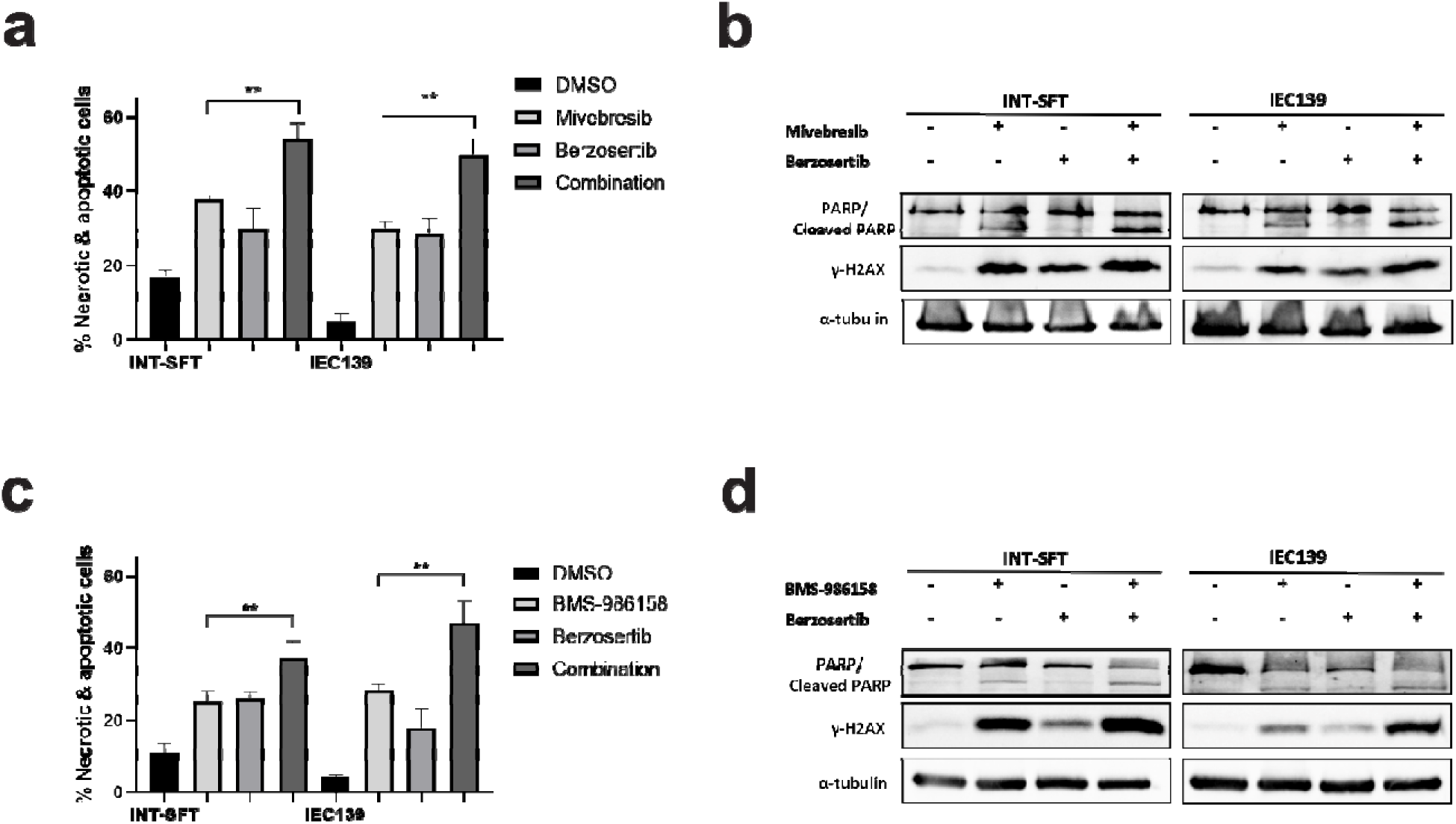
Combinatorial effects between BET inhibitors (Mivebresib and BMS-986158) and ATRi Berzosertib. **a)** Flow-cytometry-based apoptosis assays showed that combining Mivebresib and Berzosertib increased apoptotic and necrotic cell populations in INT-SFT and IEC139 cells (n=3). **b)** Western blot assays showed that combining Mivebresib and Berzosertib increased cleaved PARP-1 and γ-H2AX protein levels in INT-SFT and IEC139 cells after 72-hour treatment (n=3). The α-tubulin was used as a loading control. **c)** Flow-cytometry-based apoptosis assays showed that combining BMS-986158 and Berzosertib increased apoptotic and necrotic cell populations in INT-SFT and IEC139 cells (n=3). **d)** Western blot assays showed that combining BMS-986158 and Berzosertib increased cleaved PARP-1 and γ-H2AX protein levels in INT-SFT and IEC139 cells after 72-hour treatment (n=3). The α-tubulin was used as a loading control. For statistical analysis, two-tailed t-tests were conducted. ** denotes p < 0.01.

### *In vivo* anti-tumor efficacy testing of Mivebresib against SFTs

To evaluate the *in vivo* efficacies of Mivebresib against SFT, an IEC139 patient-derived xenograft (PDX) mouse model was used. Briefly, once IEC139 PDX tumors reach a volume of 150–200 mm³, Mivebresib (dosage: 1 mg/kg body weight, frequency: 5 consecutive days followed by a 2-day break) was administered by oral gavage (Figure 6a). As shown in Figures 6b and 6c, the treatment of Mivebresib potently reduced the tumor volumes compared to the DMSO control. More specifically, 15 days post-treatment, the tumor volumes were 270.0 ± 66.4 mm³ and 509.5 ± 60.9 mm³ for Mivebresib-treated and DMSO-treated groups, respectively. Similarly, 24 days post-treatment, Mivebresib reduced the tumor size by 65.2% (434.8 ± 178.2 mm³ vs 1247.9 ± 169.5 mm³ for Mivebresib-treated and DMSO-treated groups, respectively). It should also be noted that no significant differences in body weight were observed between Mivebresib- and DMSO-treated groups (Figure 6d), which implied that Mivebresib was generally tolerated *in vivo* at the adopted dosage. In conclusion, our data showed that Mivebresib can exert anti-tumor effects in SFT PDX models.

**Figure 6.**
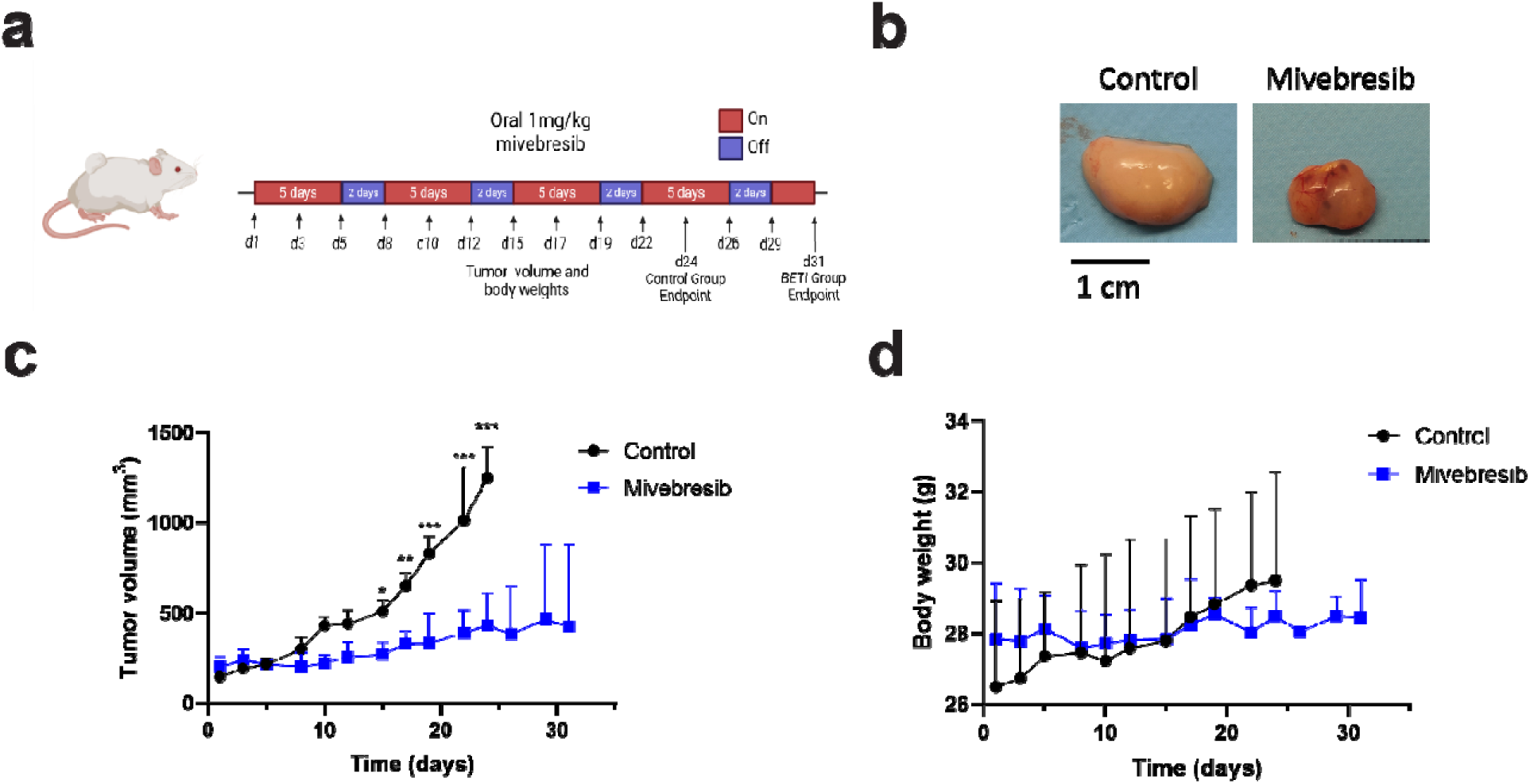
Evaluation of *in vivo* anti-tumor effects of Mivebresib in IEC139 PDX models. **a)** Schematic illustration of Mivebresib dosing regimen in IEC139 PDX models. b) Representative images of xenografts harvested at the end of the treatment period. (left) Control, (right) Mivebresib. **c)** Tumor volume progression throughout the treatment period for both Control- and Mivebresib-treated groups (n=3-4 per group). **d)** Body weight measurements throughout the treatment period for Control- and Mivebresib-treated groups (n=3-4 per group). Statistical analysis was conducted using two-way ANOVA with multiple daily comparisons (Sidak). * denotes p < 0.05; ** denotes p < 0.01; *** denotes p < 0.001; non-significant results are not shown.

## Discussion

Our study revealed that two pan-BETis, Mivebresib and BMS-986158, can potently suppress SFT cell proliferation in both *in vitro* and *in vivo* studies. These results imply that simultaneous inhibition of both bromodomains may be necessary for optimal anti-tumor effects in SFT cells. The combinatorial effects of GSK778 (BD1-selective inhibitor)^32^ and ABBV-744 (BD2-selective inhibitor)^30^ were analyzed to test this hypothesis. As shown in Supplementary Figure S2a, in both INT-SFT and IEC139 cells, the combination treatment induced more pronounced cell apoptosis than the single-agent treatment. Specifically, in INT-SFT cells, the combination treatment increased cell death by 6.1-fold and 2.0-fold compared to the GSK778 treatment and ABBV-744 treatment, respectively. Similarly, in IEC139 cells, the combination treatment increased the non-viable cell population by 13.7-fold and 6.3-fold, compared to the GSK778 treatment and ABBV-744 treatment, respectively. Consistent with cell apoptosis analysis, Western blot showed that the combination treatment induced enhanced PARP-1 cleavage and higher expression levels of γ-H2AX (Supplementary Figure S2b), further indicating that dual bromodomain inhibitions were critical for maximizing BETi efficacy in SFT. In line with these results, BETis had demonstrated activity in several other fusion gene-related sarcomas. The BET bromodomain inhibition has been described to be active in Ewing Sarcoma, suppressing the transcriptional activity of the *EWS::FLI1* transcription factor in both *in vitro* and *in vivo* xenograft experiments.^46–48^ Likewise, mivebresib treatment was shown to be active in clear cell sarcoma, reducing the protein and mRNA levels of *EWSR1::ATF1* in a dose-dependent manner. This effect was observed at the EWSR1 promoter level and was due to the modulation of BRD4 recruitment in this region.^49^ The modulation of BRD4 expression also regulates the expression level of the *PAX3/7::FOXO1* translocation in rhabdomyosarcoma.^50^ Likewise, and as shown in Supplementary Figure S3a, following 72 hours of treatment (50 nM), Mivebresib and BMS-986158 significantly suppressed the expression of *NAB2-STAT6* fusion transcripts in SFT cells. More specifically, in INT-SFT cells, Mivebresib and BMS-986158 treatments reduced *NAB2-STAT6* expression by 65.1% and 51.2%, respectively. Similarly, in IEC139 cells, Mivebresib and BMS-986158 treatments reduced *NAB2-STAT6* expression by 62.6% and 55.7%, respectively. Interestingly, other BETis (Pelabresib, ABBV-744, PLX51107, GSK778, and GSK046), which failed to induce cell apoptosis in SFT cells, did not significantly suppress the *NAB2-STAT6* expression. Indeed, as shown in Supplementary Figure S3b, a strong positive correlation between the relative expression levels of *NAB2-STAT6* and IC_50_ values for each BETi (transformed to a logarithmic scale) can be observed (Pearson correlation coefficient = 0.88, p-value < 0.001). These results suggest that BETis are active across several sarcoma histologies, including SFT, and that BET proteins may be key epi-driver factors that regulate the expression and activity of fusion gene transcription factors, through a mechanism that could be similar to other cancers driven by fusion genes, such as acute myeloid leukemia (AML), where BETi had shown promising activity.^27,51,52^

On the other hand, our results showed that BETis Mivebresib and BMS-986158, besides modulating the expression of the NAB2-STAT6 fusion gene, suppressed the proliferation of SFT cells via inducing DNA breaks and G1 cell cycle arrest. This is evidenced by ATR phosphorylation at 8-24 hours post-treatment, indicative of single-strand break (SSB) repair activation. Concurrently, upregulation of p21 and downregulation of cyclin D1 would lead to G1 cell cycle arrest. By 72 hours, SSB accumulation results in DSBs, marked by γ-H2AX, ultimately leading to apoptosis. BET proteins are implicated in DDR pathways ^53–58^, explaining the accumulation of DSBs following BETi treatment. Given that genomic instability is a hallmark of SFT progression, BETis may exploit this vulnerability ^59^. Similar findings have been reported in synovial sarcoma (SS), where mivebresib induced G1 arrest and exhibited dependency on SS18-SSX fusion expression^60^. These results are in line with previous published work, in which it was shown that BETis induced DNA damage^61,62^, through a mechanism that could be related to the RNA polymerase II (RNAPII) stalling on the chromatin, RNA:DNA hybrids (R-loops)accumulation at sites of BET protein occupancy, and the induction of DNA damage affecting the cells in S-phase.^62,63^ The increase in DNA damage due to the inhibition of BET proteins, could justify the synergy observed with DNA damage targeting agents, such as PARP and ATR inhibitors. In this sense, it has been described that BETis attenuate the repair mechanisms of double-strand break, sensitizing with PARP inhibitors in preclinical models of pancreatic ductal adenocarcinoma and ovarian cancer.^61,64^ Similar results have been published in homologous recombination–proficient cancers for the combination of BET and PARP inhibitors.^65,66^ On the other hand, the synergy of BET and ATR inhibitors, reported in our study, has also been observed in Myc-induced lymphoma cells^67^, or melanoma.^68^ The cytotoxic effect of this combination seems to be related to several biological processes, such as the induction of apoptosis, autophagy, senescence-associated secretory pathway, and ER stress.^68^

Limitations of this study, include the limited number of preclinical models of SFT, namely for *in vivo* studies. To overcome this limitation, our team is currently working on collecting fresh tissue from patients diagnosed with SFT, to attempt to establish further PDX models. To our knowledge, the IEC139 is the only non-dedifferentiated PDX model available for SFT-related preclinical research, which reinforces the relevance of the results obtained in this unique animal model. Another limitation of our study is the lack of preclinical models with the *NAB2_exon4_::STAT6_exon2_* fusion gene, to test the activity of BETis, and ensure that the activity of these compounds is similar to all SFTs, independently of the fusion genes breakpoints. While the fusion genes do not seem to impact survival^69–71^, we cannot rule out that these breakpoints can affect the efficacy of BETis in SFT.

In conclusion, our study established BETis Mivebresib and BMS-986158 as novel anti-SFT agents. Future work should focus on identifying through omics studies the epi(genetic) profile of SFTs in response to BETis and to test the combination of mivebresib or BMS-986158 with DNA damage targeting agents, such as PARP or ATR inhibitors, in in vivo models. In this sense, future work should also on understanding the mechanisms underlying the induction of DNA damage by BETis in SFT, namely on pausing of RNAPII on the chromatin and the accumulation of R-loops. Finally, a clinical trial should be designed to address the efficacy of BETis in SFT in humans and validate the preclinical shreds of evidence described in this work.

## Supporting information

For quantitative analysis, the fractions of dead/dying cells (DRAQ7 cell count/CellTracker Deep Red cell count) were calculated for each well at all t

As shown in Supplementary Figure S2a, in both INT-SFT and IEC139 cells, the combination treatment induced more pronounced cell apoptosis than the sing

Likewise, and as shown in Supplementary Figure S3a, following 72 hours of treatment (50 nM), Mivebresib and BMS-986158 significantly suppressed the ex

247 compounds were identified using the Final Timepoint effects (Supplementary Table S1),

247 compounds were identified using the Final Timepoint effects (Supplementary Table S1),

232 were determined using the AUC effects (Supplementary Table S2)

In total, 104 compounds (Supplementary Table S3, 93 from the primary screen and 11 from the clinical applications) were procured from either Selleck C

As shown in Figure 1a and Supplementary Tables S4 and S5, using these filtering conditions,

As shown in Figure 1a and Supplementary Tables S4 and S5, using these filtering conditions,

for the final timepoint effects (Supplementary Tables S6 and S7).

for the final timepoint effects (Supplementary Tables S6 and S7).

Similarly, for the AUC effects (Supplementary Tables S8 and S9), 8 hits were identified

Similarly, for the AUC effects (Supplementary Tables S8 and S9), 8 hits were identified

Therefore, we next evaluated six additional BETis (Supplementary Table S10, BMS-986158

## Acknowledgements

We thank the laboratory members in the Bleris, Hayenga, and Martin-Broto labs for their support and discussions.

## Author contributions

Conceptualization: J.L.M.-H., D.S.M., Y.L., J.M.-B., C.A.M., H.N.H., and L.B.; Investigation: J.L.M.-H., D.S.M., Y.L., J.L.M., P.G.-P., J.T.N., S.W.; Writing: all authors contributed to manuscript writing; Supervision: J.M.-B., L.B., and H.N.H.. All authors have read and agreed to the published version of the manuscript.

## Data Availability

Data collected for this article is available in the Supplementary Materials.

## Materials and methods

### Mammalian cell culture

The HCT116 cells were acquired from the American Type Culture Collection (ATCC, catalog number: CCL-247) and maintained at 37 °C, 100% humidity, and 5% CO_2_. The cells were grown in Dulbecco’s modified Eagle’s medium (DMEM media, Invitrogen, catalog number: 11965–1181) supplemented with 10% fetal bovine serum (FBS, Invitrogen, catalog number: 26140), 0.11mM MEM non-essential amino acids (Invitrogen, catalog number: 11140–050), and 0.045 units/mL of Penicillin and 0.045 units/mL of Streptomycin (Penicillin-Streptomycin liquid, Invitrogen, catalog number: 15140). To pass the cells, the adherent culture was first washed with PBS (Dulbecco’s Phosphate Buffered Saline, Mediatech, catalog number: 21-030-CM), then trypsinized with Trypsin-EDTA (0.25% Trypsin with EDTAX4Na, Invitrogen, catalog number: 25200), and finally diluted in fresh medium. The same protocol was used for maintaining NS-poly cells, NS-11, NS-17, and NS-23 cells, except that hygromycin (200 µg/mL, Thermo Fisher Scientific, catalog number: 10687010) was included in the complete growth medium for NS-poly cells.

The hTERT-immortalized human lung fibroblast cell line (Lf) was acquired from the American Type Culture Collection (catalog number: CRL-4058) and maintained at 37 °C, 100% humidity, and 5% CO_2_. The cells were grown in Fibroblast Basal Medium (ATCC, catalog Number: PCS-201-030) supplemented with Fibroblast Growth Kit-Low serum (ATCC, catalog number: PCS-201-041), and 0.3 µg/mL of puromycin (Gibco, catalog number: A1113803). To pass the cells, the adherent culture was first washed with PBS, then trypsinized with Trypsin-EDTA for Primary Cells (ATCC, Catalog number: PCS-999-003 at 37 °C for 10 min and finally diluted in fresh medium.

The primary SFT cell line (Moffitt-ns) was harvested and isolated from Moffitt Cancer Center (MCC). Under approval by the Total Cancer Care (TCC) program at Moffitt, SFT tissue samples were resected. The INT-SFT cell was gifted from Dr. Roberta Maestro’s group at the Oncology Referral Center (Centro di Riferimento Oncologico). The SKUT-1 cells were acquired from Dr. Javier Martin-Broto’s group at Advanced Therapies and Biomarkers in Sarcomas (ATBSarc). The IEC139 and CP0024 cell lines were established from female SFT and leiomyosarcoma patients, respectively, in the laboratory of Dr. Javier Martin-Broto. Moffitt-ns, INT-SFT, IEC139, and CP0024 cells were maintained in RPMI-1640 media (Thermo Fisher Scientific, catalog number: 11-875-085) containing 10% fetal bovine serum (FBS), MEM non-essential amino acids, and penicillin/streptomycin. SKUT-1 cells were maintained in DMEM media containing 10% fetal bovine serum (FBS), MEM non-essential amino acids, and penicillin/streptomycin. Cells were authenticated from primary tumors and checked for contamination every two months.

### High-Throughput Screening (HTS)

For High-Throughput Screening (HTS), a live-cell, high-content assay was employed to measure the fraction of dead/dying cells as a function of time. Optimized number of cells (400 cells/well for NS-poly, 600 cells/well for Lf, and 900 cells/well for Moffitt-ns) were plated into 384-well plates (Greiner Bio-one, catalog number: 781091) in the complete growth medium containing two dyes: CellTracker Deep Red (1:5,000, Thermo Fisher Scientific, catalog number: C34565) and DRAQ7 (1:200, Abcam, catalog number: ab109202). The cells were incubated at 37°C, 5% CO2 incubator overnight before chemical treatment using an acoustic ejection dispensing system (Echo 655 Liquid Handler, Beckman, Inc, catalog number: 001-16080). Next, images were taken for the CellTracker Red CMTPX (Ex: 561 nm) and DRAQ7 (Ex: 647 nm) channels at 0, 24, 48, and 72 hours with a 20x/0.45 air objective using an In Cell Analyzer 6000 Cell Imaging System (GE Healthcare). Four fields of view were captured per well.

Three metrics were used in data analysis: 1) AUC effects: the area-under-the-curve (AUC) of the drug response results over all time points; 2) FinalTimepoint effects: the drug response results at the final timepoint (72 hours); 3) CTG effects: the cell viability assay (Promega, CellTiter-Glo Luminescent Cell Viability assay) results at the final timepoint (72 hours).

### CellTiter-Glo Luminescent Cell Viability Assay

The CellTiter-Glo Luminescent Cell Viability kit was purchased from Promega (catalog number: G7573). The cell viability assays were performed according to the manufacturer’s recommendations. Briefly, the CellTiter-Glo reagent was added to each well in a 384-well plate (10 µL of a 1:2 dilution in PBS with 1% Triton X-100) using the Biomek i7 Automated Workstation system (Beckman Coulter, Inc., catalog number: B87581). The plates were incubated for 10 minutes at room temperature on a shaker, and subsequently, the luminescence was measured using an EnVision multimode plate reader (Perkin-Elmer, catalog number: 2105-0010).

### MTS Cell Viability Assay

All compounds were purchased from Selleck Chemicals. MTS assays were performed according to the manufacturer’s recommendations (Promega, CellTiter 96 Aqueous One Solution Cell Proliferation Assay, catalog number: G3582). Briefly, ∼5,000 cells were seeded in 96-well plates with 4 replicates. 16 hours later, the drugs were added at different concentrations. 72 hours later, MTS assays were performed by replacing the growth medium with a fresh medium containing 20% MTS substrate, and absorbances were measured at 570 nm using a plate reader (Heales, catalog number: MB-580). For data analysis, GraphPad Prism 8 was used to calculate IC_50_ values.

### Flow cytometry-based apoptosis and Cell Cycle Analysis

SFT cells were treated with candidate compounds or control DMSO. Next, for apoptosis analysis, the FITC Annexin V/propidium iodide (PI) Apoptosis Detection Kit (Immunostep) was used, following the manufacturer’s protocol using a FACSCanto II or BD Accuri C6 Plus system (BD Biosciences). The FITC fluorescence channel was used to detect annexin V-positive cells (early apoptosis), and the PerCP channel was used to detect PI-positive cells (late apoptosis/necrosis). It should be noted that a sub-G1 population was identified in INT-SFT cells, which likely indicates an apoptotic subset with degraded DNA.

For cell cycle analysis, cells were treated with candidate compounds for 24 hours and then fixed with 70% ethanol for 1 hour at 4°C. After RNase A digestion, cells were stained with PI and analyzed for DNA content using the PE channel on the BD Accuri C6 Plus system. Data was then processed with BD FACS Diva and Floreada.io software.

### *In Vivo* Treatment with Mivebresib

Nude athymic mice were implanted with 2-3 mm³ IEC139 PDX (patient-derived xenograft) tumor fragments. A minimum of 3 animals were necessary to detect a clinically relevant difference of 400mm^3^ in tumor volume, between the control-treated group and the mivebresib-treated group. The test was performed with a power of 80% and a statistical significance of 5%. The standard deviation used in the test was 125mm^3^. Once tumors reached a volume of 150–200 mm³, the mice were randomly assigned to receive either vehicle or Mivebresib (Selleckchem). Mivebresib was prepared in 2% DMSO and administered by oral gavage at a dose of 1 mg/kg body weight. The treatment followed an intermittent schedule, with dosing for 5 consecutive days followed by a 2-day break over 31 days or until tumor volume reached 1,500 mm³. Body weight and tumor volume were monitored every 2-3 days. Tumor measurements followed the same method as described in our previous report.

### Western blot (WB)

Cells were lysed using 1XRIPA buffer (1 M Tris–HCl pH 8, 0.5 M EDTA, Triton™ X-100, 10% sodium deoxycholate, 10% SDS, and 3 M NaCl), supplemented with protease and phosphatase inhibitors (Sigma-Aldrich). Protein samples (20 μg) were separated by SDS-PAGE using a constant current of 90 V for stacking and 120 V for resolving acrylamide gels. Proteins were transferred to 0.2 μm pore-size Amersham nitrocellulose membranes (Cytiva) at 4°C for 150 minutes at 200 mA constant current. Membranes were then blocked for 1 hour with 5% bovine serum albumin (BSA) or non-fat milk (PanReac AppliChem ITW Reagents) in 1X TBS 0.1% Tween-20 (Bio-Rad). Primary antibodies were then applied overnight at 4°C in BSA or milk, as recommended by the manufacturer. Next, after washing with 1X TBS-T, membranes were incubated with secondary antibodies: Rabbit Anti-Mouse IgG–Peroxidase (Sigma-Aldrich) or Goat Anti-Rabbit IgG H&L Peroxidase-conjugated (Abcam). Chemiluminescent detection was performed using ECL Prime (Cytiva), and images were acquired with a Chemidoc Imaging System (Bio-Rad). Band intensities were quantified using Image Lab software (Bio-Rad).

| Target Protein | Reference | Supplier | Host species | Blocking agent | Dilution | Molecular weight (kDa) |
| --- | --- | --- | --- | --- | --- | --- |
| **α-tubulin** | T9026 | Sigma-Aldrich | Mouse | Milk | 0.73611111 | 50 |
| **PARP-1** | 51-6639GR | BD Biosciences | Mouse | BSA | 0.5625 | 116/89 |
| **γ-H2AX (S-139)** | 9718 | Cell signaling | Rabbit | Milk | 0.73611111 | 15 |
| **p-ATR (S-428)** | 720107 | Thermo Fisher Scientific | Rabbit | Milk | 0.5625 | 300 |
| **Cyclin D1** | ab16663 | Abcam | Rabbit | Milk | 0.5625 | 36 |
| **p21** | 2947S | Cell signaling | Rabbit | Milk | 0.73611111 | 21 |
| **Wee-1** | sc-9037 | Santa Cruz Biotechnology | Rabbit | BSA | 0.5625 | 95 |
| **RAD51** | PA5-27195 | Invitrogen | Rabbit | Milk | 0.73611111 | 37 |

### Real-Time RT-PCR (Reverse Transcription-PCR)

For real-time RT-PCR assays, total RNAs were extracted using RNeasy Mini Kit (Qiagen, catalog number: 74106). First-strand cDNAs were synthesized using a QuantiTect Reverse Transcription kit (500 ng RNA, Qiagen, catalog number: 205311). Next, quantitative PCR was performed using the KAPA SYBR FAST universal qPCR Kit (Kapa Biosystems, Wilmington, MA, USA, catalog number KK4601), with GAPDH as the internal control. The forward primer for GAPDH was 51-AATCCCATCACCATCTTCCA-31, and the reverse primer for GAPDH was 51-TGGACTCCACGACGTACTCA-31. The forward primer for NAB2-STAT6 was 51-CGAAGCCACCTCTCGCAG-31, and the reverse primer for NAB2-STAT6 was 51-CTTGTAGTGGCTCCGGAAAG-31. Quantitative analysis was performed using the 2−ΔΔCt method. Fold-change values were reported as means with standard deviations.

