## Supplementary figures and images for "Identification of BET Inhibitors (BETi) Against Solitary Fibrous Tumor (SFT) Through High-Throughput Screening (HTS)"

### As shown in Supplementary Figure S2a, in both INT-SFT and IEC139 cells, the combination treatment induced more pronounced cell apoptosis than the sing

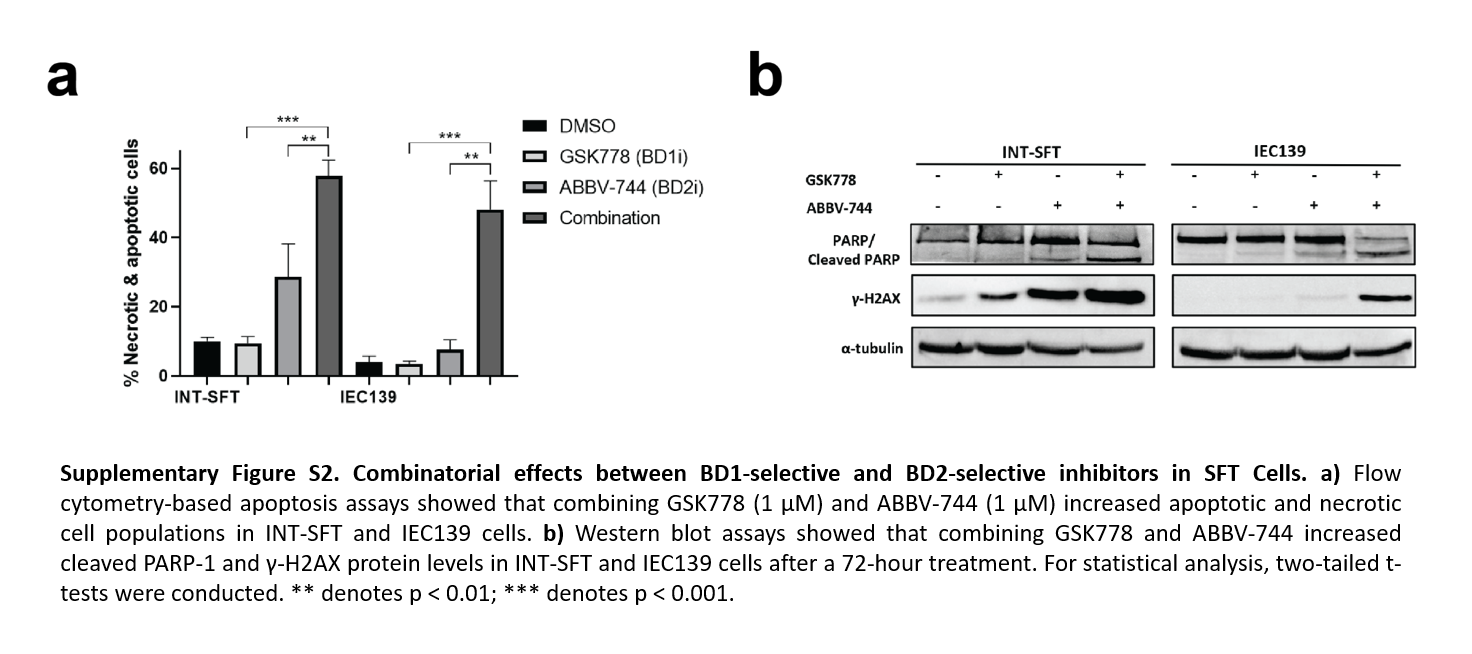

### For quantitative analysis, the fractions of dead/dying cells (DRAQ7 cell count/CellTracker Deep Red cell count) were calculated for each well at all t

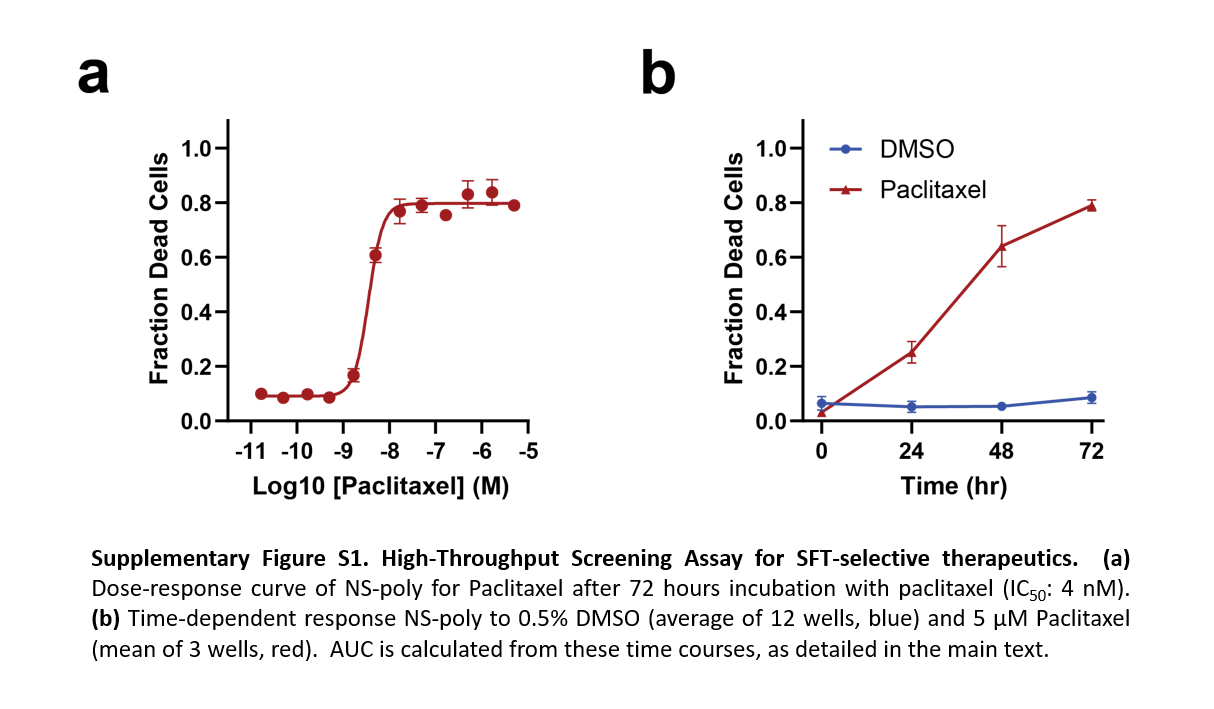

### Likewise, and as shown in Supplementary Figure S3a, following 72 hours of treatment (50 nM), Mivebresib and BMS-986158 significantly suppressed the ex

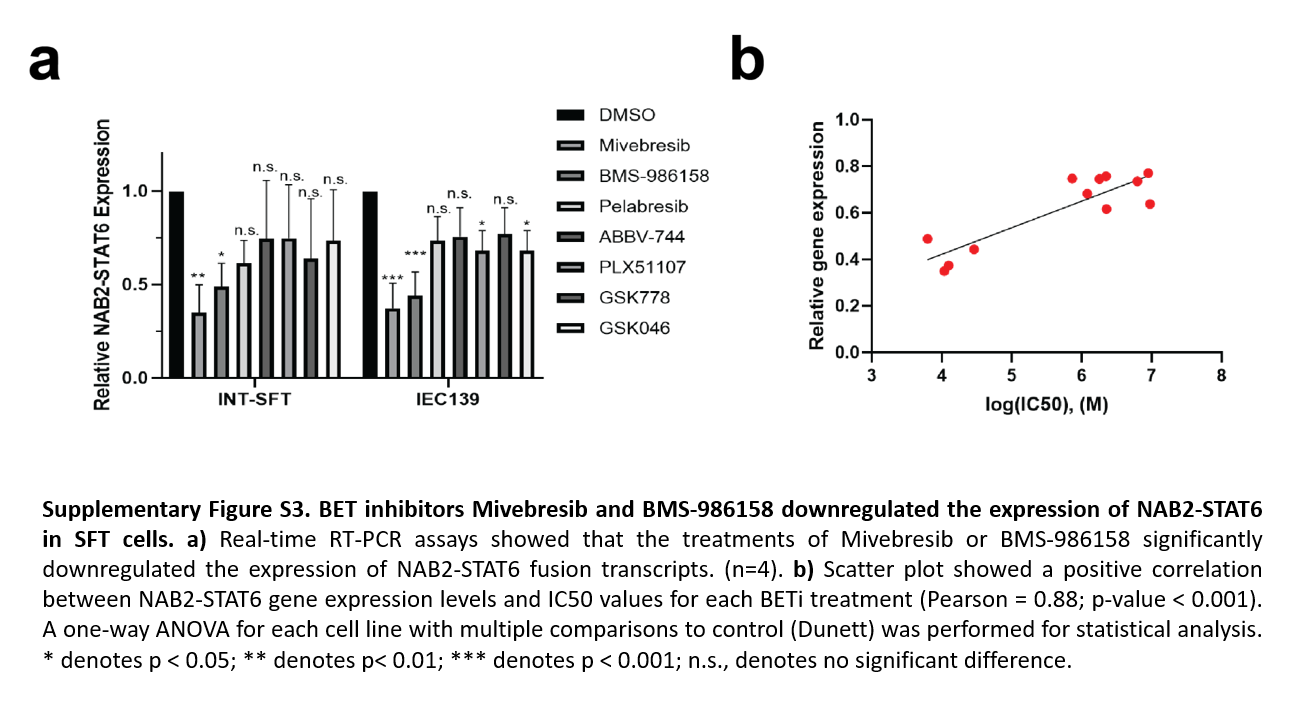
