## Supplementary material for "Identification of BET Inhibitors (BETi) Against Solitary Fibrous Tumor (SFT) Through High-Throughput Screening (HTS)": 247 compounds were identified using the Final Timepoint effects (Supplementary Table S1),

**Supplemental materials**

**Supplementary Table S1. Candidate compounds identified from the primary high-throughput screening (HTS) using the final timepoint effects.**

**Supplementary Table S2. Candidate compounds identified from the primary high-throughput screening (HTS) using the AUC effects.**

**Supplementary Table S3. Candidate compounds for the secondary high-throughput screening (HTS).**

**Supplementary Table S4. CTG effects of candidate compounds procured from NIH in the secondary high-throughput screening (HTS).** The candidate compounds are highlighted in blue.

**Supplementary Table S5. CTG effects of candidate compounds procured from Selleck Chemicals in the secondary high-throughput screening (HTS).** The candidate compounds are highlighted in blue.

**Supplementary Table S6. Final Timepoint effects of candidate compounds procured from NIH in the secondary high-throughput screening (HTS).** The candidate compounds are highlighted in blue.

**Supplementary Table S7. Final Timepoint effects of candidate compounds procured from Selleck Chemicals in the secondary high-throughput screening (HTS).** The candidate compounds are highlighted in blue.

**Supplementary Table S8. AUC effects of candidate compounds procured from NIH in secondary high-throughput screening (HTS).** The candidate compounds are highlighted in blue.

**Supplementary Table S9. AUC effects of candidate compounds procured from Selleck Chemicals in the secondary high-throughput screening (HTS).** The candidate compounds are highlighted in blue.

**Supplementary Table S10. IC50 values for additional BET inhibitors in SFT and LMS cell models.** N/A: not applicable.
